# Engineering a genetic system for female-biased, genetically unmodified progeny in mice without affecting litter size

**DOI:** 10.1101/2023.11.21.568055

**Authors:** Ido Yosef, Tridib Mahata, Xuefeng Xie, Yuhuang Chen, Hadas Bar-Joseph, Qing-Yuan Sun, Ruth Shalgi, Ariel Munitz, Motti Gerlic, Udi Qimron

## Abstract

The ability to influence the sex ratio of mammalian offspring has applications in agriculture and animal welfare. Here we describe a genetic system in mice that produces predominantly female progeny, reaching approximately ninety percent, without affecting litter size or introducing genetic modifications into the offspring. The system consists of a doxycycline-regulated cassette inserted into the Y chromosome, encoding dCas9 and RNA guides intended to impair the function of Y bearing sperm. While the cassette was originally intended to repress a spermatid maturation gene, our analyses did not detect meaningful changes in gene expression or sperm function that would account for the phenotype. Nevertheless, male mice carrying the cassette consistently produced female biased litters, and the effect was reversed by doxycycline, confirming that it depends on cassette expression. Similar female bias was observed following in vitro fertilization using sperm from transgenic males. These findings demonstrate a reproducible, genetically confined, and conditionally regulated system for sex ratio bias in mice. The strategy may prove adaptable across species, even without a complete mechanistic understanding.

**Graphical Abstract:** A single line genetic system producing female biased, genetically unmodified progeny in mice by conditionally impairing Y bearing sperm.

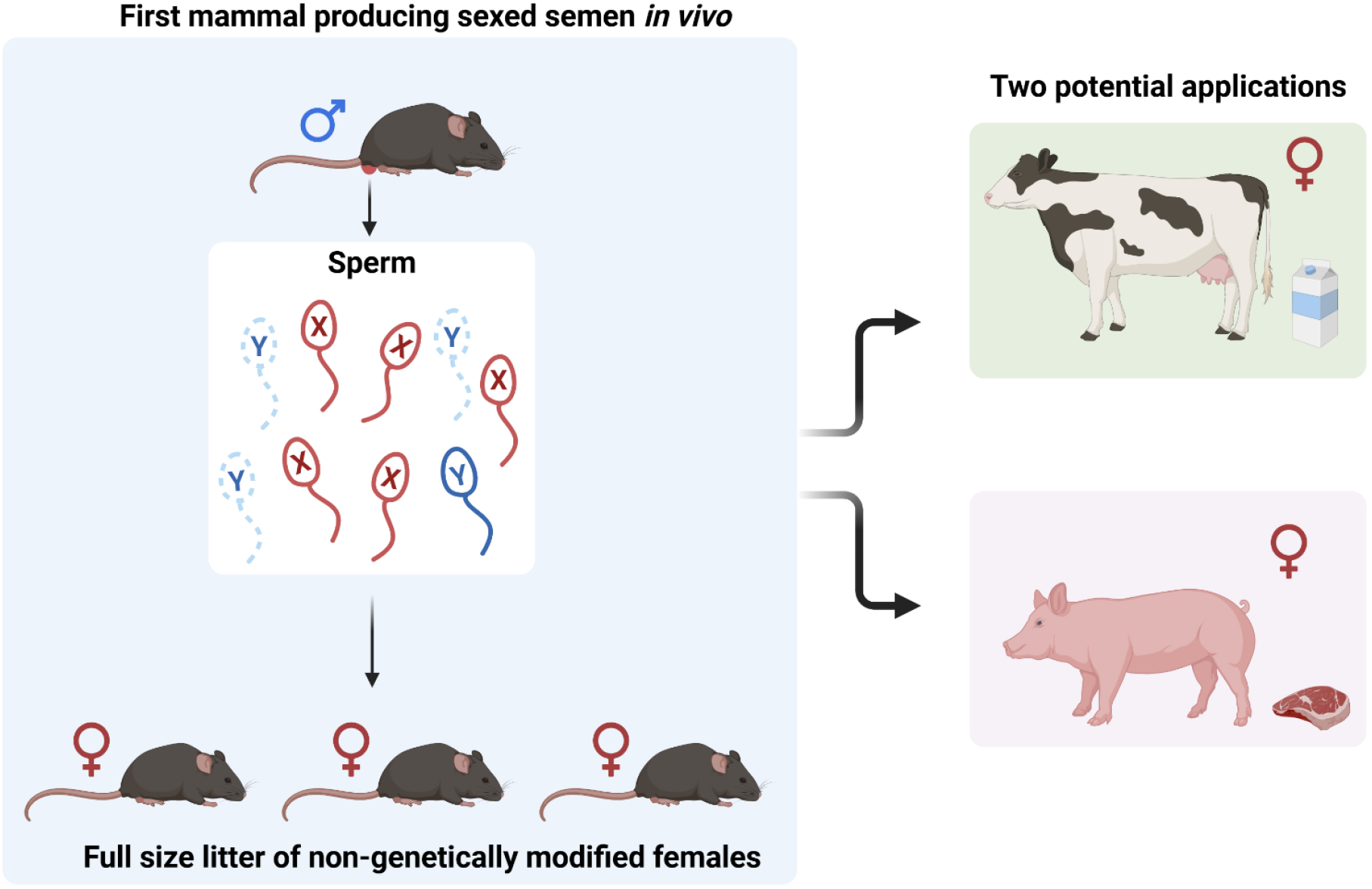

**One sentence summary:** Y chromosome manipulation yields mostly female mice, preserving litter size without genetic alteration in females.

**Significance statement:** This study describes a genetic approach for producing predominantly female offspring in mice without altering litter size or transmitting genetic modifications to the progeny. A cassette inserted into the Y chromosome of males induces a strong female bias in offspring through a mechanism that remains unclear. The female offspring do not inherit the transgene and are genetically unmodified. This system provides a simple and conditional platform for sex ratio control in mammals, with potential applications in animal breeding and welfare.

## Introduction

The ability to influence the sex ratio of mammalian offspring holds practical and ethical value in agriculture, biomedical research, and animal welfare. In livestock production, for example, female animals are often preferred for milk or egg yield, whereas in other cases, such as meat production or breeding efficiency, males may be favored. Current practices to address this mismatch between desired and actual sex ratios often involve the early culling of animals of the undesired sex, leading to economic inefficiencies and ethical concerns [1]. In laboratory animal breeding, similar inefficiencies arise when experiments require sex-matched cohorts but produce litters with unbalanced sex distributions.

Various technologies have been developed to bias sex ratios. In mammals, the most widely applied is sexed semen, where X- and Y-bearing sperm are separated by flow cytometry prior to artificial insemination [2,3]. While this method can increase the proportion of offspring of the desired sex, it is technically complex, reduces fertility, and is not suitable for all species [4–6]. Hormonal treatments, which have proven effective in aquatic species for producing single-sex populations [7–11], are not viable in mammals. Genetic strategies have been successfully implemented in insects using CRISPR-based systems or gene drives [12–20], but similar approaches in mammals remain technically challenging and often involve transgene inheritance.

We previously described a two-line genetic system in mice that used maternal expression of Cas9 and paternal inheritance of Y-linked guide RNAs to selectively eliminate male embryos post-fertilization [21]. The fundamental principles elucidated in our work were subsequently refined and replicated by the Turner group, achieving sex bias for both males and females [22]. While effective in biasing the sex ratio, those systems resulted in reduced litter sizes, genetically modified progeny, and required maintaining two genetically modified lines. A simpler and more efficient approach, particularly one that avoids transgene inheritance by the offspring and maintains litter size, would represent a valuable advance.

In this study, we designed a one-line, conditionally regulated genetic system in which a cassette encoding dCas9 and guide RNAs aimed at targeting a spermatid maturation gene was inserted into the Y chromosome. The system was intended to impair the function of Y-bearing sperm following meiosis, thereby skewing offspring sex ratios toward females. The cassette was placed under control of a tetracycline-responsive element, enabling reversible suppression of its expression. We found that male mice carrying this Y-linked cassette produced female-biased litters, with normal litter sizes and no detectable inheritance of the cassette by the progeny. The sex bias was reversed by administration of doxycycline, confirming that the effect was dependent on cassette expression.

Although the system was designed to function by repressing the targeted gene, our expression and morphological analyses did not reveal changes of sufficient magnitude to account for the phenotype. These findings suggest that the observed sex bias may result from a yet unidentified mechanism, possibly involving subtle impairments in Y-bearing sperm function. This system provides a platform for sex ratio control in mammals that is operationally simple, non-heritable, and conditionally regulated. The work highlights a reproducible and conditionally regulated system with potential applications in breeding and animal welfare.

## Results

### Generation of transgenic male mice harboring a conditional repressive cassette on the Y chromosome

To generate male mice that would predominantly produce X-bearing sperm, we designed a genetic cassette encoding dCas9 [23] fused to a KRAB repressor domain, along with five RNA guides targeting the *Spem1* gene. This gene is known to be required for proper maturation of spermatids in mice [24]. The entire cassette was placed under the control of a tetracycline-responsive element (TRE), allowing repression of dCas9 expression in the presence of doxycycline [25]. The cassette was flanked by homology arms targeting an intron within the Y-linked *Uty* gene [26], previously shown to be suitable for insertion of genetic elements [21]. The design rationale was that after meiotic segregation, only the Y-bearing gametes would express the cassette and undergo repression of *Spem1*, while X-bearing gametes would be unaffected. As a result, we expected the Y-bearing sperm to be nonfunctional or less competitive, leading to a female-biased progeny (Figure 1).

**Figure 1.**
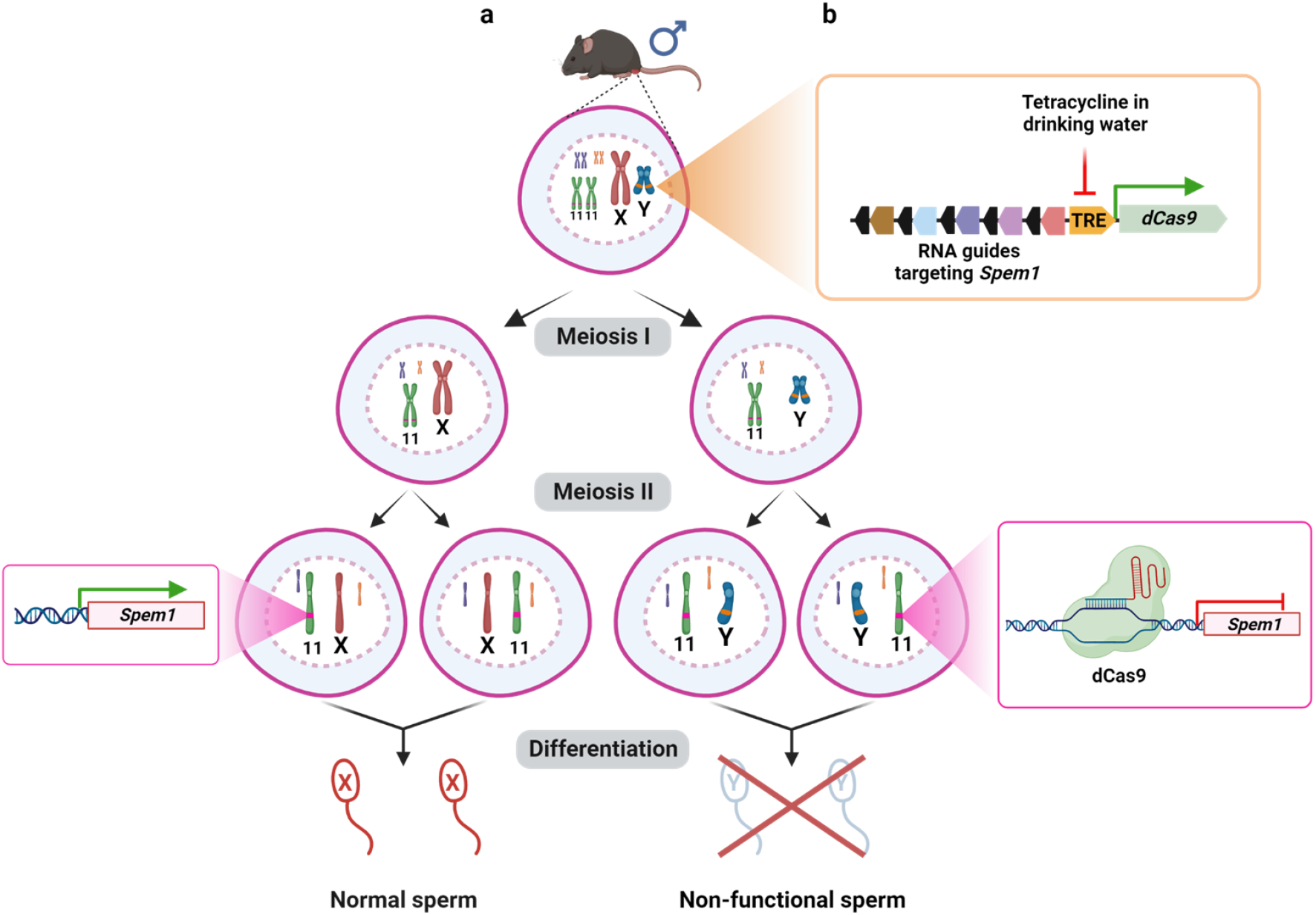
Sexing the semen *in vivo* in a male mouse. **(a)** A male mouse’s Y chromosome is genetically engineered to contain a cassette intended to repress a gene essential for gamete development. After meiosis, the Y-gametes encode and express the cassette, whereas the X-gametes lack it. As designed, the Y-gametes are expected to fail to mature while the X-gametes develop normally. **(b)** A schematic representation of the cassette inserted into the Y chromosome. dCas9 is expressed from a promoter encoding a tetracycline-responsive element (TRE), repressed if tetracycline/doxycycline is present in the drinking water. The cassette also encodes five RNA guides that target the *Spem1* gene encoded on chromosome 11. The representation is not drawn to scale. Additional elements of the cassette are described in Materials and Methods.

The cassette was introduced into C57BL/6 mouse embryonic stem cells through homologous recombination, and properly targeted clones were used to generate chimeric mice [27]. The resulting F_0_ transgenic males exhibited poor mating efficiency, which necessitated the use of in vitro fertilization (IVF) to establish the line. All male founders carried the transgene on their Y chromosome, as validated by PCR (Figure S1), and were otherwise viable and morphologically normal. Male offspring derived from IVF were backcrossed to wild type females to generate F_1_ and subsequently F_2_ transgenic males, which were used for the breeding experiments described below. These progeny mice could breed normally. One consistent difference was a reduction of approximately 25% in body weight relative to wild type mice at 8 weeks of age (Table S1), though this phenotype had not been previously associated with *Spem1* knockout [24]. Nevertheless, this was not expected to compromise the strategy, as the male mice are used solely for breeding, and the objective is to produce predominantly female offspring. Aside from the weight difference, the transgenic mice developed normally, with no physiological or morphological abnormalities (Figure S2a). In this regard, it is important to note that the female offspring of the transgenic mice maintained normal body weight, as determined by comparing their growth curve to the published reference values for wild-type C57BL/6 females (Figure S2b).

### Breeding experiments reveal highly female-biased litters from transgenic males

To test whether the transgene could bias the sex ratio as designed, we carried out a series of controlled matings. We first mated six wild type C57BL/6N males with two wild type females each, resulting in a total of 61 pups. As expected, the sex distribution in this group was close to even, with 43% males and 57% females (Figure 2a). We next mated 15 transgenic males, each with two wild type C57BL/6N females. These matings yielded 157 pups in total, with a striking female bias of 89% females and 11% males. All male progeny in this group were positive for the Y-linked transgene, whereas all female pups tested negative, as determined by PCR (Figure S1). Importantly, the average litter size in this group was not significantly different from that of wild type controls (Figure 2b), indicating that the sex bias was not the result of selective embryonic loss.

**Figure 2.**
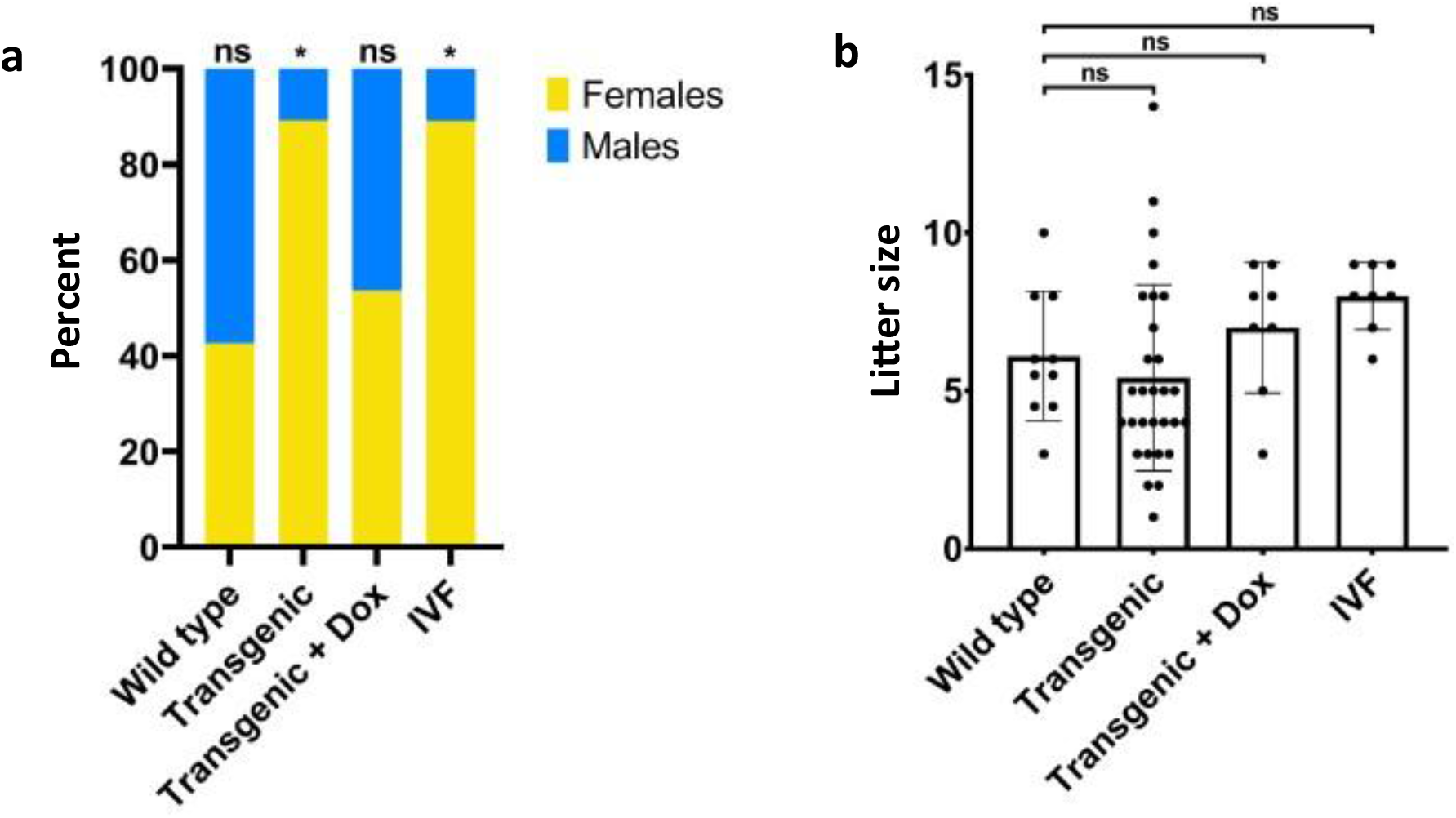
(**a**) The sex distribution of the total pups from crosses between C57BL/6N females with wild type C57BL/6N males (n=61), F_2_ transgenic males (n=157), F_2_ transgenic males treated with 2mg/ml doxycycline (Transgenic + dox) (n=56), and F_2_ transgenic sperm through IVF (n=64). (**b**) Litter sizes of the crosses indicated in panel a. Bars represent average litter size ± Standard Deviation. Significance was determined by using unpaired parametric two-tailed t-test. *, *P* < 0.0001; ns, not significant.

To test whether the observed bias was indeed dependent on expression of the cassette, we administered doxycycline into the drinking water of a separate group of eight transgenic males prior and throughout the mating period. These animals were housed individually with two females each, and the resulting 56 pups displayed a sex ratio of 46% males and 54% females, a result statistically indistinguishable from wild type controls (Figure 2). This observation supports the conclusion that cassette activity is responsible for the sex bias and that the effect can be conditionally turned off by doxycycline.

### Female bias is preserved in vitro and not associated with inheritance of the transgene

To assess whether the observed sex bias could be recapitulated in vitro, we performed IVF using sperm isolated from four transgenic males. These sperm were used to inseminate oocytes collected from wild type females. The fertilized zygotes were transferred to pseudopregnant recipient females, and a total of 64 offspring were obtained. Of these, 89% were female, a distribution nearly identical to that observed *in vivo* (Figure 2). As with the *in vivo* litters, none of the female pups carried the Y-linked cassette. Thus, the cassette reliably biases offspring toward females without being inherited by the progeny.

### *Spem1* expression and sperm phenotype do not fully explain the observed effect

Although the phenotype was statistically robust and reproducible, we were surprised to find that *Spem1* expression in the testes of transgenic males was only slightly and insignificantly reduced compared to wild type, as determined by quantitative PCR (Figure S3). Since X-bearing and Y-bearing gametes coexist in the same tissue, and *Spem1* is expressed late in spermatid maturation, we considered the possibility that transcripts from X-bearing cells may mask the reduction in Y-bearing gametes.

We further analyzed the morphology and function of sperm from three transgenic males, three transgenic males treated with doxycycline, and three wild type controls. The testes of transgenic mice, whether treated or untreated with doxycycline, were morphologically similar to those of the control group (Figure S4a) but were smaller even after adjusting for overall weight differences (Figure S4b). Importantly, despite the size difference, sperm count per epididymis remained comparable across all three groups (Figure S4c). The testes and the cauda epididymis appeared similar in histological staining (Figure S5), though a few of the seminiferous tubules in the transgenic mice (either treated or not with doxycycline) appeared abnormal. Computer assisted sperm analyses for the sperm of the three groups indicated no difference in direct motility, progressive motility, path velocity, progressive velocity, or track speed for sperm from all groups (Figure S6). Furthermore, Y-chromosome probe labeling of the causa epididymal sperm, showed equal proportions of X- and Y-bearing sperm in the transgenic and wild type groups (Figure S7). These findings rule out skewed gamete production or major defects in motility as causes for the observed bias.

Morphological analysis did reveal a specific difference: a higher proportion of sperm in transgenic mice exhibited a tightly bent head structure (8.9% ± 1.5%), compared to 3.8% ± 0.9% in wild type and 4.5% ± 1.8% in doxycycline-treated transgenic mice (Figure 3). This morphological defect has been previously associated with *Spem1* deficiency [24], but the frequency of affected sperm was far below the expected 50% if all Y-bearing gametes were impaired. Thus, while the morphology data are compatible with partial impairment of Y-bearing sperm, they do not explain the striking magnitude of the observed female bias.

**Figure 3.**
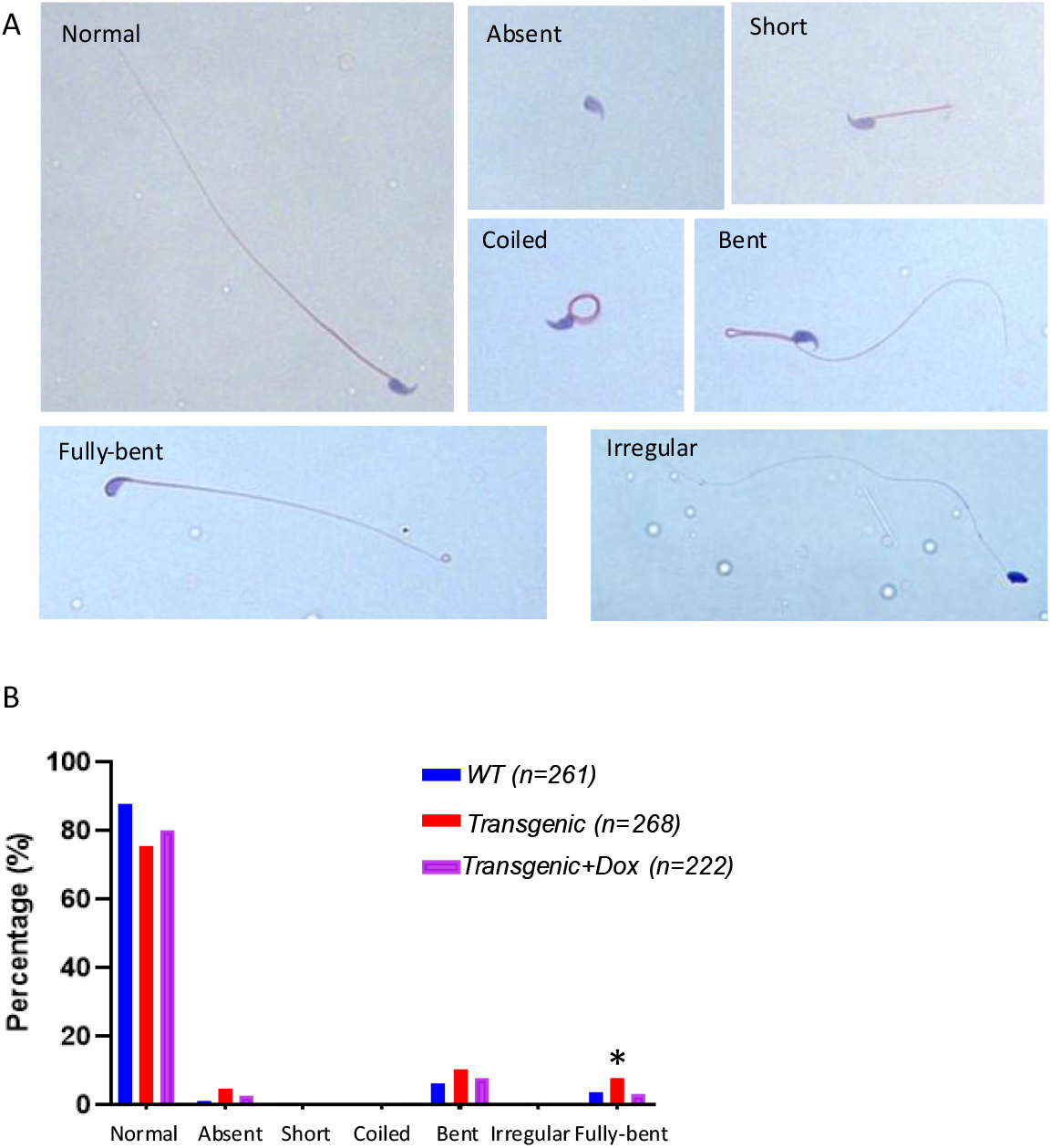
Sperm morphology of WT mice, transgenic mice, and transgenic mice treated with doxycycline. A. Representative images of sperm from the cauda epididymides of different mice showing the indicated morphologies. B. Distribution of the different morphologies in the different groups. *, P<0.05

## Discussion

In this study, we established a genetic system in mice that biases offspring toward females without reducing litter size and without transmitting any genetic modification to the progeny. This outcome was achieved by expressing a repressive cassette from the Y chromosome of male mice, though the precise mechanism by which this cassette acts remains to be elucidated. The cassette was designed to impair the maturation of Y-bearing gametes by targeting a gene known to be required for spermatid differentiation. The system was conditionally regulated through a tetracycline-responsive promoter, allowing for suppression of cassette expression by doxycycline administration.

The transgenic males produced strongly female-biased litters, with up to 89% of offspring being female, while maintaining normal fertility and average litter sizes. The bias was reversed by providing doxycycline in the drinking water, supporting the conclusion that the cassette is directly responsible for the effect. Importantly, female offspring did not inherit the transgene, confirming that the cassette is not transmitted through the X-bearing sperm. These findings demonstrate that the sex bias is linked to the presence and expression of the cassette on the Y chromosome.

We initially reasoned that the cassette would function by repressing *Spem1*, a gene required for proper spermatid maturation [24]. However, our measurements showed only a slight and statistically insignificant reduction in *Spem1* expression in the testes of transgenic males. Morphological analysis of sperm revealed a small but significant increase in abnormal sperm with bent head morphology, previously associated with *Spem1* deletion [24]. Nonetheless, the frequency of these defects was low and not sufficient to account for the magnitude of the observed female bias. Sperm motility, gamete counts, and the ratio of X- and Y-bearing gametes were not significantly altered. Thus, although the phenotype is reproducible and cassette-dependent, the mechanism underlying the selective impairment of Y-bearing sperm remains unclear.

While the initial design of the system relied on the hypothesis that repression of *Spem1* would impair Y-bearing sperm, our findings suggest that the observed phenotype is unlikely to be mediated by substantial reduction in *Spem1* levels. One possibility is that the cassette affects a tightly regulated or spatially confined process during spermiogenesis, in a manner that is not detectable by our standard assays. Alternatively, the burden of expressing dCas9 and guides from the Y chromosome itself may compromise Y-bearing sperm in a subtle but effective way. It is also possible that unintended interactions between the expressed RNA guides and off-target genomic regions contribute to the phenotype. Future studies will be required to dissect whether epigenetic, transcriptomic, or structural perturbations contribute to the observed sex bias. The ability to generate sex-biased litters without compromising fertility or transmitting transgenes opens the possibility of adapting similar systems to other mammalian species. In principle, a comparable strategy could be constructed by identifying a late-stage expression gene required for spermatogenesis in the target species and placing a repressive cassette on the Y chromosome under conditional control. The choice of target gene and promoter would require species-specific validation, but it would be of interest to test whether similar constructs can produce equivalent outcomes in other species. Such systems may find future applications in agriculture, where skewing the sex ratio toward the more economically relevant sex could improve efficiency and reduce the need for postnatal sex sorting.

## Materials and Methods

### Targeting vector for transgene insertion

The targeting vector VB201210-1271-bpw was synthesized by VectorBuilder (USA). It includes the following elements: Uty homology arms (a 4.6-kb 5′ arm and 2.5-kb 3′ arm) for insertion into intron 2 of Uty; cassette of 5 sgRNA against *Spem1*; Kruppel-associated box (KRAB) domain fused to dead Cas9 with a nuclear localization signal (KRAB-dCas9-NLS); Self-deleting Neomycine (SDN) cassette; tetracycline-controlled transactivator (tTA); a negative selection marker - diphtheria toxin A (DTA) cassette downstream of the 3′ homology arm. The full sequence and annotations of the vector can be found here: https://en.vectorbuilder.com/vector/VB201210-1271bpw.html.

### Housing and husbandry

All mice were housed and bred under specific pathogen-free conditions maintained in the Cyagen Transgenic Animal Center. Animals were housed in exhaust ventilated cage systems or individually ventilated cage systems. Air in the facility was controlled by fully enclosed ventilation systems. Temperature was kept at 24±2^0^C. Humidity was kept at 40-70%. Light was automatically set from 6AM to 6PM. Pre-sterilized diet was manufactured according to the standard published by the Chinese government (No. GB 14924.3-2010, Laboratory Animals: Nutrients for Formula Feeds). The quality of diet was monitored via periodic testing by state-approved third-party test providers. Water was filtered from the city’s public water system, placed in bottles with sipper tubes and autoclaved before use. The bedding was placed in cages and autoclaved before use. Water was added with either 0.5 mg/ml or 2mg/ml Doxycycline hyclate (TCI, catalog number D4116) where indicated and replaced weekly. Mating was carried out by placing one male with two female mice in a cage with nesting materials. The weight of the males was monitored weekly. Copulatory plugs, pregnancies, and births were monitored every three days.

### Transgenic mice

Transgenic mice were produced by Cyagen Biosciences Inc., Suzhou, China. The linearized targeting vector VB201210-1271-bpw was introduced into Cyagen’s proprietary TurboKnockout embryonic stem (ES) cells from C57BL/6N by electroporation. Clones that underwent homologous recombination were isolated using positive (neomycin) and negative (DTA) selection. 96 G418 resistant clones were picked and amplified in 96-well plates. A duplicate of this 96-well plate served for DNA isolation and subsequent PCR screening for homologous recombination. Of the 41 initially targeted ES clones, 18 were randomly selected for recovery and expansion. From these, 5 clones showing good cell morphology were selected, frozen, and subjected to Southern blot and karyotype analysis to confirm correct targeting and chromosomal integrity. Six targeted ES clones were injected into pre-blastocyst-stage albino B6N embryos and transplanted into CD-1 pseudo pregnant mice. This procedure yielded 24 F_0_ mice with 100% chimerism and 3 F_0_ mice with approximately 90% chimerism, as judged by coat color. Three independent F_0_-derived lines were established, and male founders were selected randomly. Males were genotyped as described below for validating transgene insertion and germline transmission.

### Sperm extraction and analysis

Four-month-old male mice were euthanized by cervical dislocation. Following previously described methods (46), epididymides were harvested and minced in Dulbecco’s Modified Eagle Medium (DMEM). The tissue fragments were incubated for 30 minutes at 37°C in a humidified incubator to allow sperm release. Sperm concentration was determined using a hemocytometer.

For morphological analysis, sperm smears were prepared on slides, fixed, and stained with H&E (Phygene, PH0516) according to the manufacturer’s instructions. Morphological abnormalities were assessed by examining at least 200 spermatozoa per mouse.

To evaluate sperm motility, multiple incisions were made in the epididymis, which was then placed in human tubal fluid supplemented with 10% FBS and incubated at 37°C for 5 minutes. Sperm motility parameters were analyzed using Computer Assisted Sperm Analysis (CASA).

### Genotyping and sex determination

Genomic DNA templates for PCR reactions were extracted from mice tails. The transgene and sex of the pups were determined by 3 PCRs of the Y chromosome using oligonucleotides described in Table S2 and LongAmp DNA polymerase (New England Biolabs, USA). Thermocycling conditions were as follows: 94 °C (3 min), 34 X cycles of 95 °C (30 s), 60 °C (30 s), 65 °C (50 s/kb) followed by 72 °C (3 min) and 4 °C (hold). Sex was further confirmed at day 7 and at weaning by observing the genitals.

### *In vitro* fertilization

Sperm was collected from different transgenic mice and stored in liquid nitrogen. Oocytes were obtained from female mice 15-17 hours following chorionic gonadotropin injection. The fertilization dish containing the cumulus-oocytes complexes (COCs) was incubated on CARD medium for 30-60 min. before insemination. 20 μL of the sperm was thawed, diluted in FERTIUP sperm dish and incubated for 30 min. 10-20 μL of sperm was then added to the COCs and incubated for 3 h. The oocytes were then washed 3 times in a fresh mHTF medium. 6 H after insemination, oocytes that had only one pronucleus were removed. After overnight, the 2-cell stage embryos were then transferred to female mice.

## Acknowledgements

UQ is supported by the European Research Council – Horizon 2020 research and innovation program, grant no. 818878. Cody Genetics Ltd. sponsored the IVF experiments. AM received funding from the US-Israel Bi-national Science Foundation (grant no. 2015163) and the Israel Science Foundation (grant no. 542/20). MG received funding from the US-Israel Bi-national Science Foundation (grant no. 2017176) and the Israel Science Foundation (grant no. 2174/22). We thank Nitzan Gonen and Anat Eldar-Boock for critical comments and suggestions.

## Author Contributions

Conceptualization, U.Q.; Methodology, I.Y., H.B-J. R.S., A.M., M.G., and U.Q.; Investigation, I.Y., J.C., H.B-J. R.S., A.M., M.G., and U.Q.; Writing – Original Draft, U.Q.; Writing – Review & Editing, I.Y., T.M., R.S., A.M., M.G., and U.Q.; Funding Acquisition, A.M., M.G., and U.Q.

## Competing interests

I.Y., M.G., and U.Q. have submitted a patent request on a related technology in January 2019. I.Y., A.M., M.G, and U.Q. have submitted a patent request on the described technology in August 2023.

## Data Availability Statement

The data that support the findings of this study are available from the corresponding author upon request.

## Notes

### Summary of Updates

This revised version includes extensive changes in response to peer review. The manuscript has been substantially rewritten to clarify that the observed female-biased phenotype is robust and reproducible, but that the underlying mechanism remains unresolved. We have removed speculative mechanistic interpretations and explicitly reframed the Spem1-targeting strategy as a design rationale rather than a demonstrated causal pathway. We also corrected minor grammatical issues, improved figure legends to reflect design intent rather than confirmed outcomes, and streamlined the discussion to reduce redundancy and strengthen logical flow. No new data were added in this version.

