## Supplementary files for "Engineering a genetic system for female-biased, genetically unmodified progeny in mice without affecting litter size"

### Engineering mice for female-biased progeny without impacting genetic integrity and litter size

#### Supplementary Figures

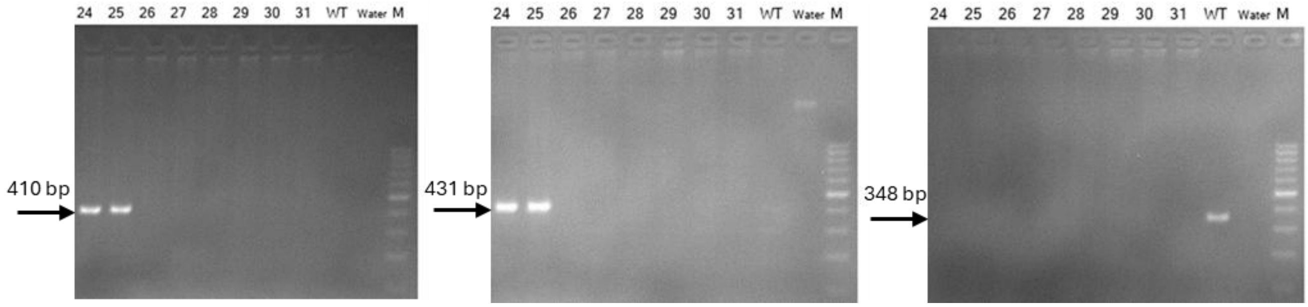


**Figure S1. Representative Gel images of products from PCRs 1 (left), 2 (middle), and 3 (right) (see Table S2 for PCR details).** Eight progeny mice and one wild-type control were tested, mice 24, 25, and the control were males and the rest were females, as also confirmed by visual examination. Gels are representative examples of over 150 PCR validations.

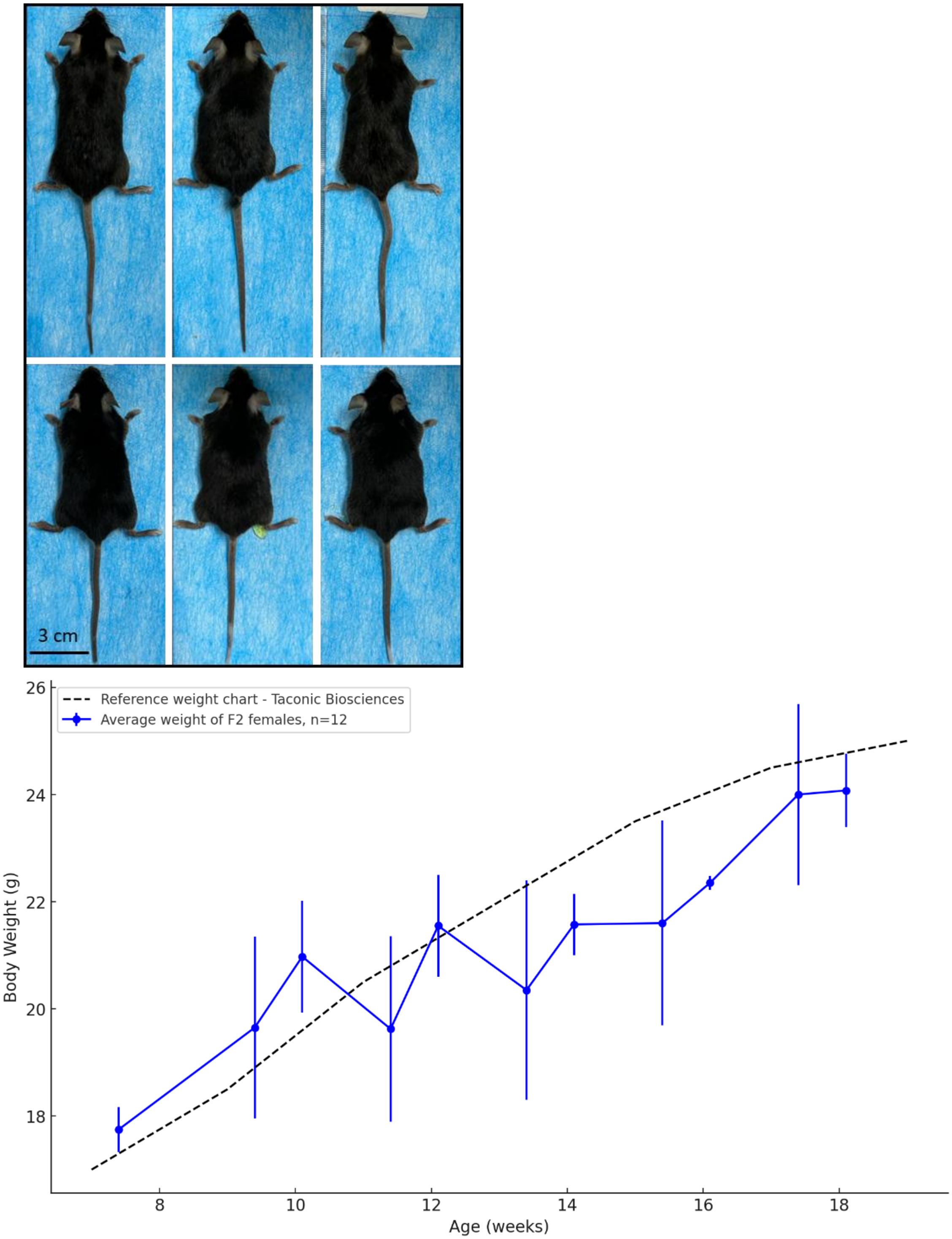


**Figure S2. A. Images of three C57BL/6N mice (top) and three transgenic mice (bottom)**. Apart from weight, no other differences were observed. **B.** Average body weight of F2 female mice (n=12) measured from 7 to 18 weeks of age, compared to the reference growth curve of C57BL/6 females (Taconic Biosciences). Error bars indicate standard deviation.

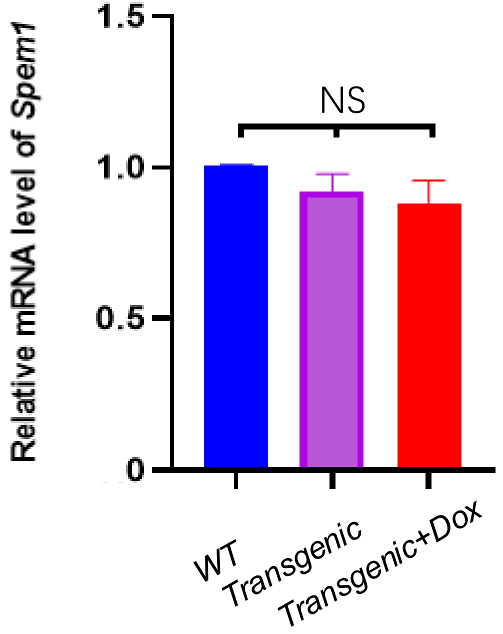


**Figure S3. Relative mRNA levels of Spem1 as determined by qRT-PCR.** Bars represent mean + standard error. n=5 for WT and Transgenic+Dox, 6 for Trangenic.

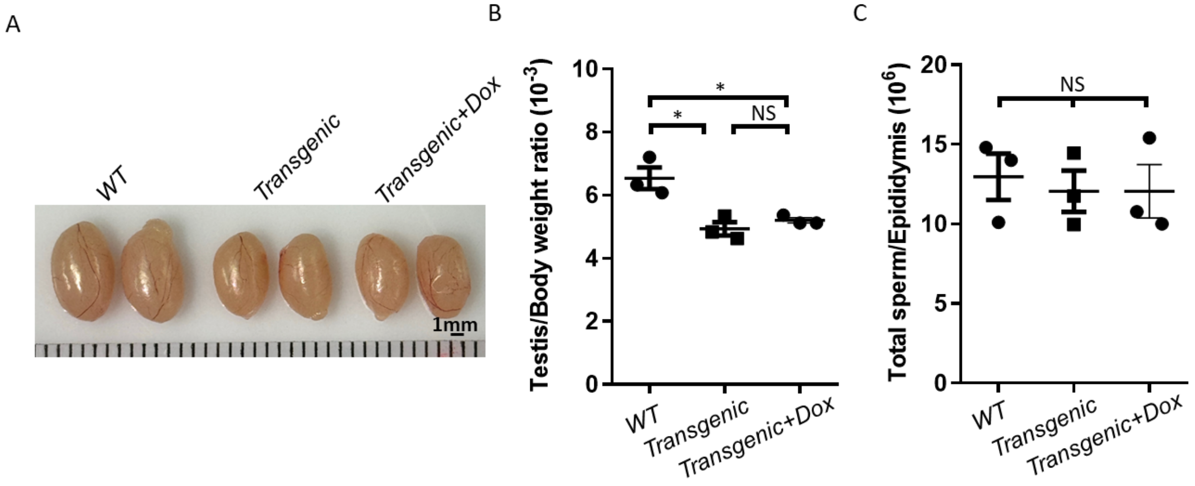


**Figure S4. Testes appearance, size, and sperm counts.** A. Representative images of testes from WT, Transgenic and Transgenic+Dox mice. B. Testes/body weight ratio of WT, Transgenic and Transgenic+Dox mice. C. Total sperm number per epididymis WT, Transgenic and Transgenic+Dox mice. *, $p<0.05$; NS, not significant.

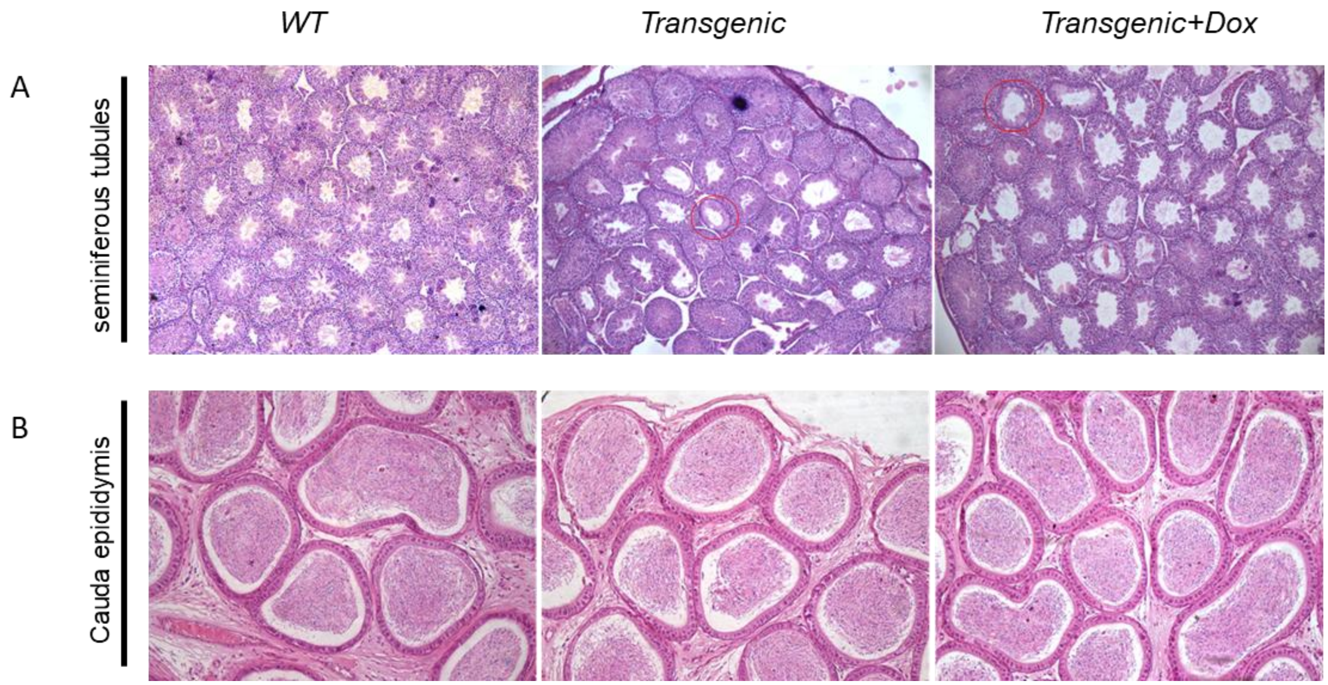


**Figure S5. Representative images of:** A. Seminiferous tubules and B. Cauda epididymis sections of WT, Transgenic and Transgenic+Dox mice after H&E staining. Abnormal seminiferous tubules are indicated by the red circles.

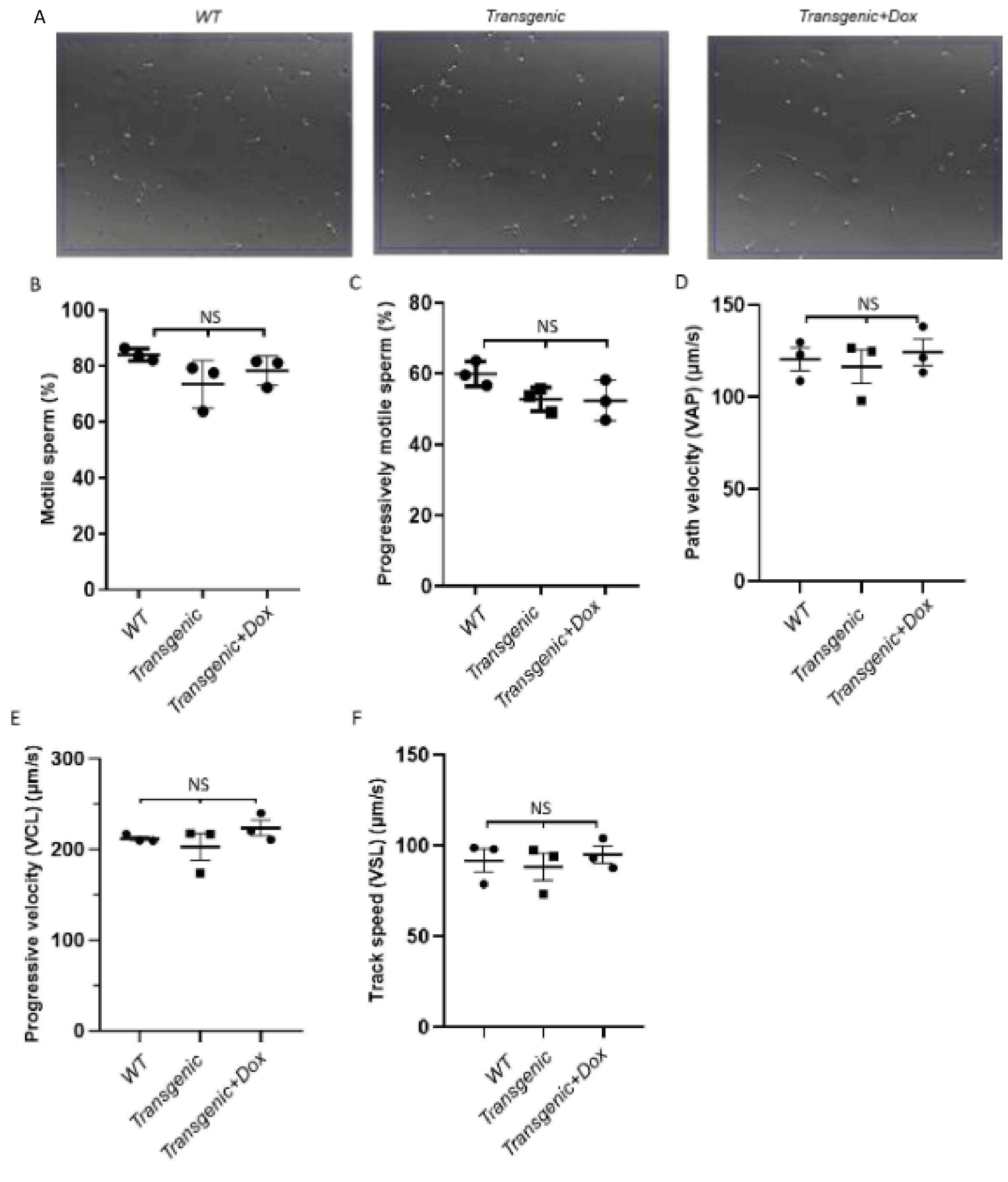


**Figure S6.** A. Representative images of motility analysis samples sperm of WT, Transgenic and Transgenic+Dox mice. B. Percentages of motile sperm from WT, Transgenic and Transgenic+Dox mice . C. Percentages of progressively motile sperm from WT, Transgenic and Transgenic+Dox mice. D. The average path velocity (VAP) of the sperm in WT, Transgenetic and Transgenetic+Dox mice. E. The average straight-line velocity (VSL) of the sperm in WT, Transgenetic and Transgenetic+Dox mice. F. The average curvilinear velocity (VCL) of the spermatozoa sperm in WT, Transgenetic and Transgenetic+Dox mice. NS, not significant.

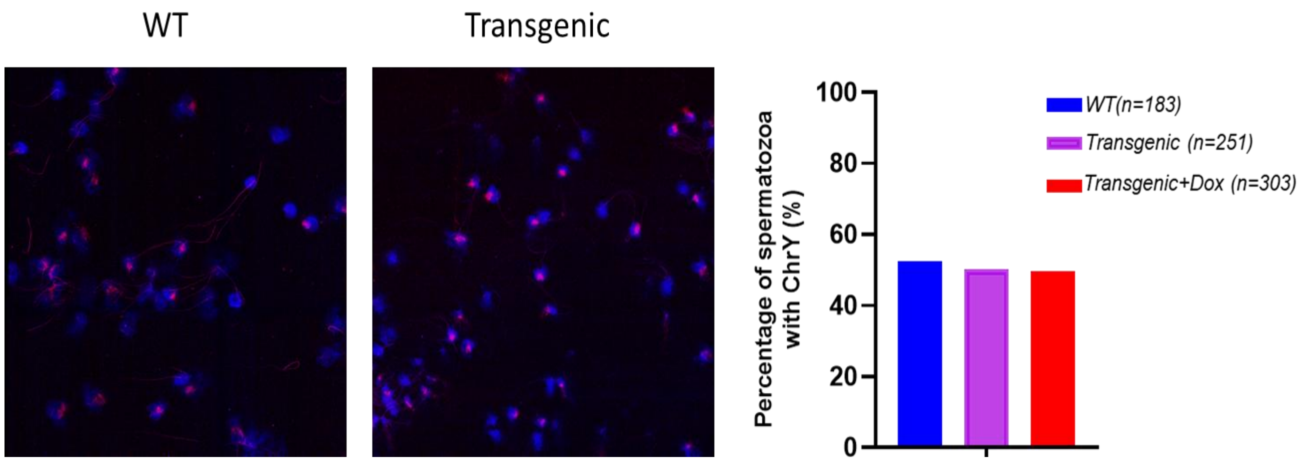


**Figure S7. Percentage of Y-gametes.** Representative images of ChrY probing in sperm of wild type and transgenic mice (2 left images) and quantification of these percentages in the three groups (right).

#### Supplementary Tables

**Table S1. Transgenic vs. WT mice body weight at ~8 weeks of age.**

| Group | Mouse ID | Weight (g) | Average (g) | Standard deviation |
|---|---|---|---|---|
| Transgenic - Dox 2mg/ml | 76 | 22.0 | 18.4 | 2.05 |
| | 79 | 18.2 | | |
| | 69 | Dead | | |
| | 64 | 17.2 | | |
| | 34 | 17.2 | | |
| | 24 | 17.4 | | |
| Transgenic - Dox 0.5mg/ml | 47 | 21.2 | 18.1 | 2.17 |
| | 10 | 17.6 | | |
| | 19 | 18.1 | | |
| | 8 | 19.3 | | |
| | 32 | 17.9 | | |
| | 80 | 14.6 | | |
| Transgenic - No treatment | 68 | 22.0 | 17.5 | 2.68 |
| | 78 | 18.7 | | |
| | 25 | 14.4 | | |
| | 35 | 17.4 | | |
| | 81 | 15.5 | | |
| | 18 | 16.8 | | |
| C57BL/6N No treatment | 1 | 23.3 | 24.2 | 1.05 |
| | 2 | 23.5 | | |
| | 3 | 24.2 | | |
| | 4 | 23.7 | | |
| | 5 | 26.2 | | |
| | 6 | 24.2 | | |

**Table S2. List of primers used in this study.**

| Primer | Sequence (5’-3’) | Comments |
|---|---|---|
| TKIE100-F1 | GGTTGTTGCAGCCTCTTGTGA | PCR 1 to confirm transgene insertion (amplifies left junction of insertion). Expected size – 410 bp. Unmodified mice – no amplification. |
| TKIE100-R1 | CGCCCCAACCTCATTGTGA | |
| TKIE100-F3 | CTTAGACATGCTCCCAGCCGA | PCR 2 to confirm transgene insertion (amplifies right junction of insertion). Expected size – 431 bp. Unmodified mice – no amplification. |
| TKIE100-R2 | AACTGTTTCATTTCCCCTCTCCT | |
| TKIE100-F4 | GGTTGTTGCAGCCTCTTGTGAT | PCR 3 to confirm transgene insertion (amplifies a region flanking the transgene). Expected size – 348 bp. Transgenic mice – no amplification (over 10kbp). |
| TKIE100-R4 | TGGTAAAACTACCCTGTTTCCTTTC | |

### Supplementary Materials and Methods

#### Quantitative PCR

Total RNA was extracted using the TransZol Up Plus RNA Kit (TransGen Biotech, Cat# ET111-01-V2,Beijing, China), followed by cDNA synthesis with the HiScript II 1st Strand cDNA Synthesis Kit (Vazyme, Cat# R211-02, Nanjing, China), both performed according to the manufacturers' protocols. Quantitative PCR analysis was conducted on a LightCycler® 480 Instrument II System (Roche, Cat# 05015278001, Basel, Switzerland) using AceQ qPCR Probe Master Mix (Vazyme, Cat# Q111-03, Nanjing, China). Gene expression levels of Spem1 were quantified using the $\Delta\Delta$Ct method with $\beta$-actin (ACTB) as the endogenous reference. Relative mRNA expression fold changes were determined by comparing experimental groups (transgenic and transgenic + dox) to the control group. The sequences of all primers (5' to -3') used are listed below.

RT-PCR primers for Spem1:

Fw: ATGGCCATGGCTGAGCGGCC

Rv: GGGTCACCAAATTGATGCCA

RT-PCR primers for actin:

Fw: AGAAGAGCTATGAGCTGCCT

Rv: TCATCGTACTCCTGCTTGCT

#### Determination of testes/body weight ratio

Before euthanasia, body weights of four-month-old mice were recorded. Following euthanasia, testes were excised and immediately weighed. The gonadosomatic index (GSI) was calculated using the formula: (testicular weight [mg]/body weight [g]) $\times 10^3$ ) to express testis mass relative to body mass.

#### Histology analysis

As described previously (1), four-month-old mice were euthanized by cervical dislocation, and their epididymides and testes were collected and fixed in Bouin's solution. The tissues were subsequently dehydrated through a graded ethanol series and embedded in paraffin. Serial sections were prepared and stained with hematoxylin and eosin (H&E). Images were captured using a Digital Leica DFC700 T camera mounted on a Leica DM2500 microscope.

**FISH (ChrY probing methodology)**

Sperm FISH analysis was performed on samples collected from control, transgenic, and transgenic+dox groups. The cauda epididymis was isolated from each group, and sperm were released and incubated at 37° C for 30 minutes in a 5% $CO_2$ atmosphere. Collected sperm were purified by centrifugation (300g, 5 minutes) in PBS to remove debris, followed by double fixation with Carnoy's solution and spreading onto slides. For sperm chromatin decondensation, samples were treated with 1N NaOH and incubated in 5mM DTT at 37° C for up to 150 seconds. The slides were then dehydrated through a graded ethanol series (70%, 85%, 100%) and heated at 80° C to ensure complete ethanol evaporation. FISH was performed using a Y chromosome-specific probe (Empire Genome, Chromosome Y Red [MCENY-10-RE]). After probe denaturation at 85° C for 10 minutes, hybridization was carried out for 24 hours at 37° C in a humidified chamber. Post-hybridization washes were performed twice with 2×SSC containing 0.1% Tween at 65° C, followed by two washes with 2×SSC at room temperature, each for 5 minutes. Finally, the slides were counterstained with DAPI.


1. X. Xie, *et al.*, Biallelic HFM1 variants cause non-obstructive azoospermia with meiotic arrest in humans by impairing crossover formation to varying degrees. *Hum Reprod* 37, 1664–1677 (2022).
